# Novel joint enrichment test demonstrates high performance in simulations and identifies cell-types with enriched expression of inflammatory bowel disease risk loci

**DOI:** 10.1101/2023.06.13.544716

**Authors:** Alexandru-Ioan Voda, Luke Jostins-Dean

## Abstract

A number of methods have been developed to assess the enrichment of polygenic risk variants – from summary statistics of genome-wide association studies (GWAS) – within specific gene-sets, pathways, or cell-type signatures. The assumptions made by these methods vary, which leads to differences in results and performance across different genetic trait architectures and sample sizes. We devise a novel statistical test that combines independent signals from each of three commonly-used enrichment tests (LDSC, MAGMA & SNPsea) into a single P-value, called the block jackknife GWAS joint enrichment test (GWASJET). Through simulations, we show that this method has comparable or greater power than competing methods across a range of sample sizes and trait architectures. We use our new test in an extensive analysis of the cell-type specific enrichment of genetic risk for inflammatory bowel disease (IBD), including Crohn’s disease (CD) and ulcerative colitis (UC). Counterintuitively, we find stronger enrichments of IBD risk genes in older gene expression data from bulk immune cell-types than in single-cell data from inflamed patient intestinal samples. We demonstrate that GWASJET removes many seemingly-spurious enriched cell-types identified by other methods, and identifies a core set of immune cells that express IBD risk genes, particularly myeloid cells that have been experimentally stimulated. We also demonstrate that many cell-types are differentially enriched for CD compared to UC risk genes, for example gamma-delta T cells show stronger enrichment for CD than UC risk genes.

**Author summary:** Genetic association studies have discovered a number of DNA variations that are associated with heritable human diseases and traits. One method of investigating the functions of these variants is to test whether they are enriched in parts of the genome associated with specific cell-types or cell conditions – defined by gene expression data or other similar data types. However, there are a number of published statistical methods to test such enrichments; these methdos make different assumptions and their results can vary, sometimes dramatically. We present a novel consensus method, called GWASJET, that combines the results of these different methods to produce a single result. We show that GWASJET can outperform individual methods in simulations. We apply this method to gene expression data from a number of tissues and conditions relevant to inflammatory bowel diseases (IBD). Our method removes potentially false results based on *a priori* biological knowledge, and reveals that IBD genes are generally clustered in a large number of immune cell-types, especially myeloid cells treated with specific stimulatory molecules.

## Introduction

Genome-wide association studies (GWAS) have been instrumental in uncovering the genetic basis of complex diseases and traits [1]. Thousands of loci have been found to be associated with a range of heritable phenotypes. However, the functions of most of these loci are not yet fully understood [2], and determining their impacts on downstream pathways will help us better understand disease processes, which in turn can significantly improve the process of developing new therapeutics [3]. Many research groups are working on scaling up post-GWAS functional experiments to understand the impact of these risk variants on gene expression and other up- and down-stream cellular phenotypes [4]. These functional experiments have produced a wide range of datasets, including traditional data such as genome-wide gene expression datasets [5], and newer datasets such as genome-wide epigenetic assays like ATAC-Seq [6] and detailed single-cell gene expression maps, including from patient samples [7]. One key type of tool used in post-GWAS studies to understand risk loci is GWAS enrichment testing. These statistical tools are designed to estimate the enrichment of risk variants in a given set of genomic regions, or a given gene-set, and test whether this enrichment is significantly higher than expected under the null hypothesis. When paired with the results of functional genomics experiments on cell or tissue samples (such as transcriptomic or epigenetic profiling), enrichment testing can be used to identify cell-types or cell-states which express disproportionate numbers of risk genes for a given disease. This approach has been successful applied to identify cell-types associated with genetic risk for many diseases [8–10]. In this paper we specifically focus on enrichment testing for cell-type specific signals, but the same approach can be used to test for enrichment of other gene-sets or genomic regions, such as canonical pathways [11]. Many GWAS enrichment testing tools have been developed and successfully applied and among them, MAGMA [12], LDSC [5,13,14] and SNPsea [15] have been quite popular. However, the modeling and assumptions made by these methods vary markedly, with LDSC applying a polygenic model across all common variants, MAGMA modelling disease risk on a per-gene basis, and SNPsea analysing only smaller numbers of genome-wide significant loci. As such, the enrichment estimates and p-values for a given dataset differ depending on which method is chosen, sometimes dramatically so. Even once a single tool is selected, there are also other choices that need to be made, such as whether to treat genomic regions or gene-sets as discrete (inside vs outside) or whether to use a continuous measure for every gene or region (e.g. a continuous cell-type specificity score). As would be predicted from these different assumptions (and as we will demonstrate through simulations in this study), different genetic architectures and study features (heritability, polygenicity, sample size, etc) can change what the best performing method is for a given study.

Moreover, there are several ways of obtaining gene-sets for enrichment testing from gene expression data: one can use canonical differential expression analyses, such as DESeq2 & edgeR [16,17]. But there are many more gene-specificity scoring algorithms being used for enrichment tests. For example, one LDSC enrichment testing study has used t-statistics to define cell-type specific gene-sets from expression data [5]. Another MAGMA enrichment study [18] has used a method called EWCE [19] to select cell-type specific genes from expression data. It is possible that the various gene selection approaches may have advantages and disadvantages under various biological and technical contexts.

Taken together, this means that researchers who wish to test for enrichment of risk variants in cell-types have to make complex choices about both the enrichment testing tool and the gene selection approach. To get around this problem, in this paper we introduce a new technique, called GWASJET, which can combine the results of different enrichment testing results into a single p-value.

We use simulations to investigate the performance of this new approach, and then use this tool to carry out a detailed study of cell-type enrichment in inflammatory bowel disease (IBD). IBD is a prime example of a complex disease where accurate GWAS enrichment testing can provide valuable insight. IBD is commonly classified by disease location and other phenotypic differences into two subtypes, Crohn’s disease (CD) and ulcerative colitis (UC). They are both characterised by chronic gastrointestinal inflammation in genetically-predisposed individuals upon exposure to environmental risk factors. IBD is severe (even deadly without treatment) and it affects hundreds of thousands of people in the United Kingdom alone [20].

Earlier GWAS enrichment tests with IBD data showed a significant enrichment of risk loci within genes that are specific to immune and gastrointestinal cells, including T cells and myeloid cells, as well as epithelial cells [5,8,21]. New gastrointestinal RNA-sequencing data, including single-cell data from patient intestinal biopsies, have recently become available. These can enable us to characterise rarer cell populations, compare more conditions (location, treatments, etc.) and examine differences between cell-types from patients and controls. For these purposes, projects such as the Human Cell Atlas [22] may yield novel findings in IBD, especially when combined with a more accurate joint test for GWAS enrichment.

This paper has three aims: firstly, to describe GWASJET, including a new R package that implements it; secondly, to conduct GWAS simulations to assess the performance of the joint test and its individual components across a range of plausible use cases; thirdly, to apply the novel test to a broad range of cell-types from real datasets to identify those that can be prioritized in inflammatory bowel disease.

## Results

### Power across a range of enrichment strengths, GWAS sample sizes and trait heritabilities

With the approach described in the methods section, we can simulate GWAS at desired levels of enrichment, heritability, etc.

Assessing the type 1 error rate using the null simulations (a causal density ratio of 1, i.e. a heritable trait with no enrichment of signal within the gene-set), we found that all enrichment methods are slightly conservative (Figure 1A), as they produced less than the expected 5% rate of false positives (our method producing a false positive 2% of the time).

**Figure 1:**
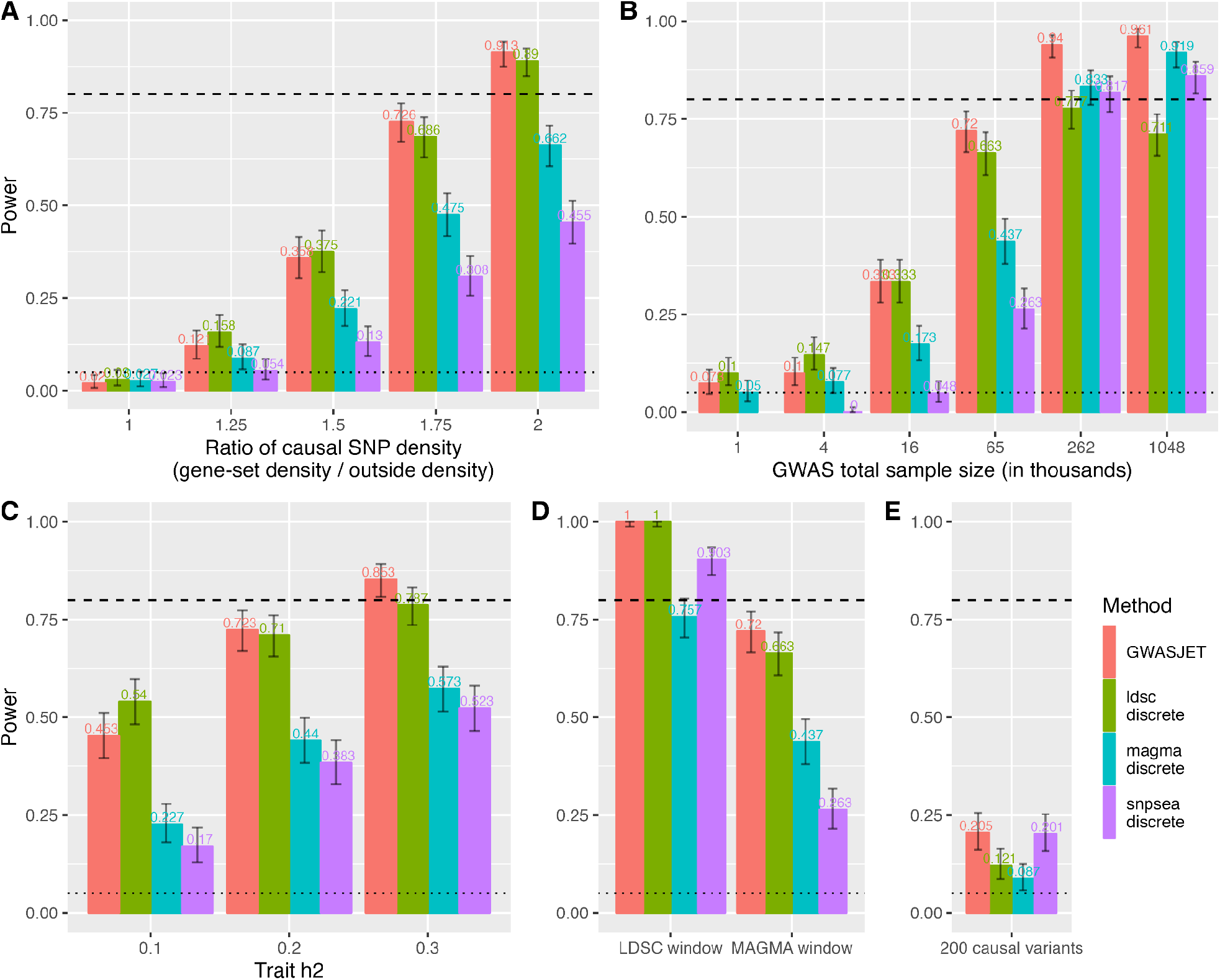
GWAS simulations to compare the statistical power of MAGMA, LDSC, SNPsea and GWASJET. **A)** The power to detect causal gene-sets with increasing heritability, which is controlled in these simulations by changing the ratio of causal SNP densities in the gene-set region versus outside regions. A ratio of 1 represents the null hypothesis (no enrichment of heritability in the gene-set). **B)** Increasing GWAS sample size, from 1 thousand to 1 million individuals, leads to increases in statistical power. **C)** The effect of trait heritability on power. **D)** The influence of the SNP-to-gene mapping window (100kb as described in LDSC model paper, or 10kb as described in MAGMA’s paper). The ratio of causal SNP density in the MAGMA window is set as before (to 1.75), but the causal density ratio in the LDSC window is decreased (set to 1.25) to adjust for the increase of linked-SNPs due to the larger window. **E)** A less polygenic scenario, where only 200 causal variants are sampled per simulation, instead of 4,000 as in the other simulations. In all plots power is determined at p < 0.05, and error bars are 95% confidence intervals on the estimated power.

As expected, power increases for all methods when gene-sets hold a higher proportion of heritability (fig. 1A), when GWAS are larger (fig. 1B), or the trait is more heritable (fig. 1C). Starting with a comparison of previously published methods, we observe that LDSC had higher power than MAGMA across most scenarios, until GWAS sample size reaches biobank scales (fig. 1B), above 200 thousand cases and controls, at which point LDSC’s power began to saturate and MAGMA overtook it. In general, SNPsea had lower power across almost all scenarios, except for a slight advantage in an oligogenic model (fig 1E). Interestingly, simulating GWAS under the MAGMA proximity window (for SNP-to-gene mapping) does not necessarily confer an advantage to MAGMA when compared to simulating under LDSC’s proximity window simulation parameters (fig. 1D).

GWASJET retains or exceeds the power of the leading method in all simulated contexts. In almost all scenarios the difference between the best single test and GWASJET was small, but for any given test there were scenarios where GWASJET beat it by a substantial margin. For example, GWASJET substantially outperforms LDSC but not MAGMA and SNPsea at very large sample sizes, substantially outperforms MAGMA and SNPsea but not LDSC for low heritability traits and outperforms LDSC and MAGMA but not SNPsea for less polygenic traits.

We next assessed the computational footprint of each method, measured in CPU hours on a single core with different numbers of gene-sets (Table 1). MAGMA and SNPsea have a relatively small computational cost, and MAGMA in particular scales well with the number of gene-sets tested as most of its computation is in pre-processing the summary statistics. LDSC is much slower, mostly due to the need to calculate partitioned LD scores for each gene-set, and GWASJET has a much larger computational footprint than any of the other methods, as the single threaded runtime is a high multiple of the MAGMA, LDSC and SNPsea runtimes. Roughly 75% of GWASJET’s runtime (for 10 gene-sets) is spent computing with SNPsea 201 times, which, unlike MAGMA and LDSC, has no significant precomputing stage, and thus has to be run in full in every block. A further ∼15% is spent running LDSC, of which almost the entire time is spent calculating LD scores.

**Table 1:** Method runtimes for a single-core. Tests are carried on a single core of an Intel Xeon™ 6126 CPU (2.6 GHz) with RAM access capped to maximum 15 GB. The joint test is many times slower than LDSC, MAGMA and SNPsea. Any pre-processing steps in SNPsea are negligible, with the analysis permutations dominating essentially all of SNPsea’s computational time. MAGMA and LDSC’s running time are broken down into their two main computational steps.

| # of gene-sets tested | MAGMA |  |  | LDSC |  |  | SNPsea | GWASJET |
| --- | --- | --- | --- | --- | --- | --- | --- | --- |
|  | Gene Z scoring | Final regression | Total | LD scoring | Final regression | Total | Total | Total |
| 1 | 11m0s | 14.5 sec | 11m15s | 1h39m | 36 sec | 1h40m | 1m13s | 8h47m |
| 10 |  | 14.9 sec | 11m15s | 8h12m | 1m16s | 8h14m | 12m21s | 54h52m |

We applied our novel GWASJET method to find cell-types enriched for IBD risk across a range of IBD-relevant tissues. In terms of broad cell-type enrichment, our results reproduce what is already known about IBD risk: that it is strongly enriched in a host of immune cells, including myeloid cells, T cells and NK cells (Fig 2A).

**Figure 2:**
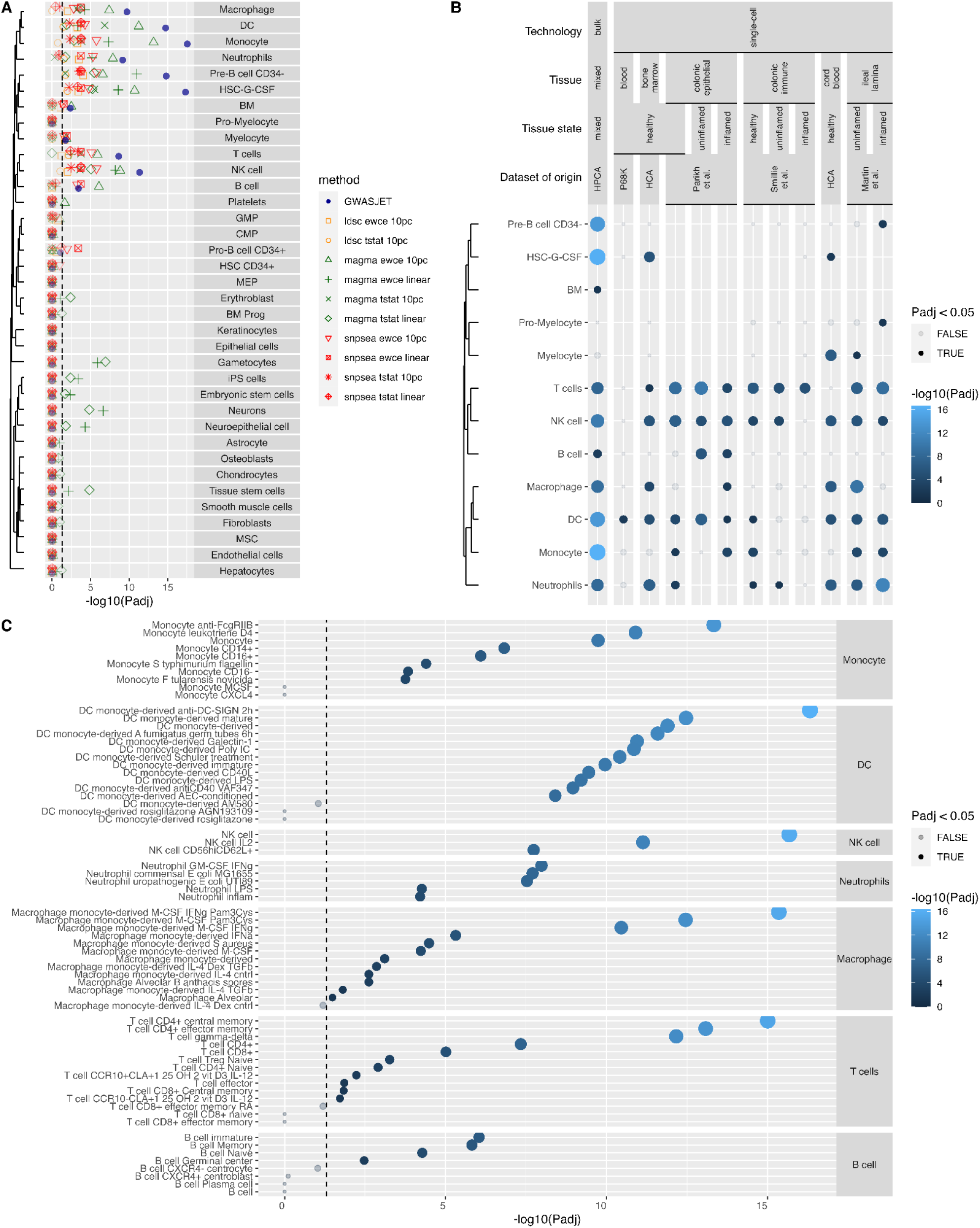
Joint enrichment test for IBD loci in expression data from broad and fine-grained types of cells. **A)** Results from the block jackknife joint GWAS enrichment test and its 10 component tests for IBD loci within broad cell-types from the Human Primary Cell Atlas. In the legend, “10pc” refers to testing the top 10% specifically expressed genes, while “linear” stands for testing all genes using a continuous cell specificity score, ewce and tstat represent two different cell specificity scoring methods and GWASJET, ldsc, magma and snpsea represent different enrichment testing methods. The dashed line shows adjusted p-value < 0.05. **B)** Joint enrichment test results for IBD loci across different expression datasets, broken down by technology (single-cell vs bulk), tissue type and disease state (healthy, uninflamed IBD tissue, inflamed IBD tissue). **C)** Enrichments in fine-grained cell subtypes of significant enriched broad cell-types from the HPCA analysis.

**Figure 3:**
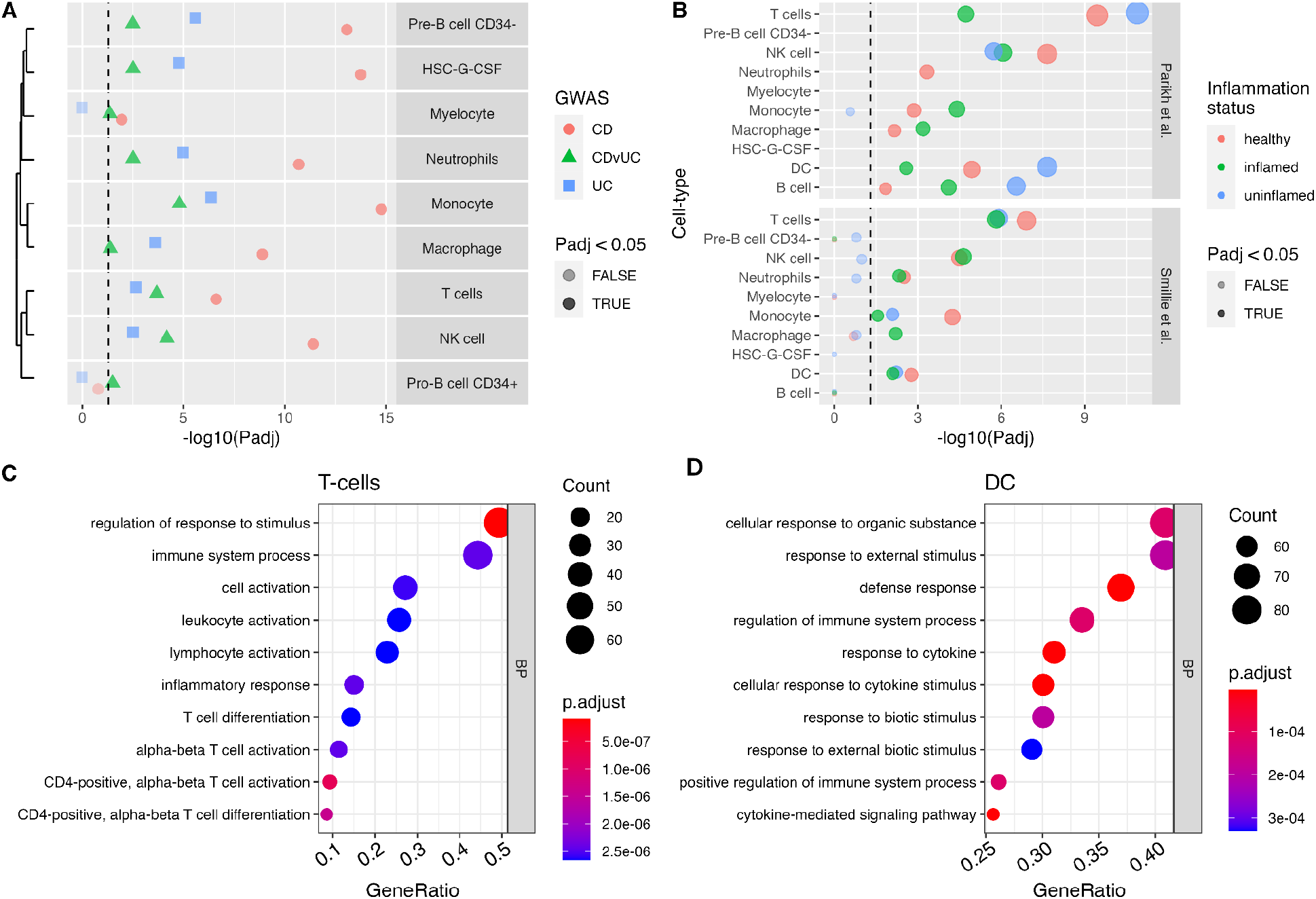
IBD GWAS enrichment in cell-types across disease subtypes, inflammation status and cell-type specific risk pathways. A) Enrichment of risk loci for cell-type specific gene expression for CD and UC, along with enrichment for loci that differ between CD and UC (CDvUC), showing only the significant broad cell-types from HPCA. The dashed line shows adjusted p < 0.05. B) Enrichment of IBD loci within cell-types taken from tissue with different inflammation statuses, using data from two single-cell studies of IBD patient and control samples C & D) Enrichment of GO terms within cell-type specific IBD risk genes for two cell broad cell-types (T cells and Dendritic Cells), compared to cell-type specific genes for this cell-type that are not risk genes. All classes of GO term were tested, but all significant results were BP (Biological Process) terms.

For sufficiently strong enrichment results, most enrichment testing methods produce the same answer. For instance, all 10 of the component tests found a significant enrichment of IBD risk in dendritic cells. However, an unlucky choice of enrichment test can miss very important well established findings - for instance, unlike the other methods used, the recommended choice of settings for LDSC (top 10% mode with t-statistic gene selection) does not find a significant enrichment of IBD risk in monocytes (Padj=0.18). GWASJET assigns a very low p-value to this cell-type (Padj=2.9e-18). Some methods also produce statistically significant but seemingly spurious enrichment results, for instance both MAGMA linear modes find a highly significant (Padj < 1e-5) enrichment of IBD risk in gametocytes. Because GWASJET takes a consensus of methods, these sporadic outliers driven from a single method are more effectively controlled for.

### Application of joint GWAS enrichment test to inflammatory bowel disease

Looking across tissues and disease states (fig 1B), we saw that lymphocyte and myeloid cell-types tended to maintain enrichment in expression of IBD risk genes across tissues and conditions. Soberingly, the strongest enrichments were observed in the (decade old, microarray-based, bulk) human primary cell atlas, which showed more significant results for almost all enriched immune cell-types than newer, single-cell, patient tissue datasets.

The high power and low false positive rate of GWASJET makes it well suited for testing for enrichment within finer-grained cell-types (fig 2C). We saw the greatest enrichment within T cell subsets in memory CD4 T cells and gamma-delta T cells, and within B cells enrichment was driven by naive and memory B cells (with no signal coming from plasma cells). Among the myeloid cells, the strongest enrichment in all cases was seen in some form of stimulated, rather than resting or ex-vivo, cells. The opposite effect was also seen, where certain cell stimulations removed the enrichment of IBD GWAS signal, for instance, unlike almost all monocyte subsets, monocytes treated with the rosiglitazone, a synthetic PPARγ activator [23], showed no enrichment in IBD signal (Padj = 1).

One understudied area in the genetics of IBD is the difference between the genetics of CD and UC. We derived new post-hoc summary statistics for a CD vs UC GWAS and tested for cell-types that were enriched for risk genes that varied between these two diseases. We found that most of the major immune cell-types differed between CD and UC to some extent, and that in all cases the enrichment for risk within the cell-type was stronger in CD than in UC (fig 3A). The largest CD vs UC enrichment at the broad cell-type was in monocytes, though at the finer-grained cell-type level (Supplementary Figure 1) we saw an even stronger difference between CD and UC was seen for gamma-delta T cells.

Given the recent appearance of multiple single-cell gene expression datasets from inflamed patient tissue, it stands to reason that immune cells within these datasets would show a greater enrichment of IBD risk variants. However, we did not find this to be the case in general when we examined two studies with both health and inflamed tissue (fig 3B). In some cases, for instance in macrophages, we did observe greater enrichment in inflamed tissue. However, most others showed stronger enrichment in non-inflamed tissue in both studies (e.g. T cells, dendritic cells), or mixed results in the two studies (e.g. monocytes, NK cells).

Finally, we tested the functions IBD risk genes played in enriched cell-types, by focusing on differences in gene ontology (GO) terms between IBD-associated cell-type-specific and non-IBD associated cell-type-specific genes. We found that cell-type-specific IBD risk genes tend to affect core pathways of the immune cells they are enriched in, such as activation and differentiation in T-cells (fig 3C) or responses to cytokines or other stimulations in dendritic cells (fig 3D).

## Discussion

This study introduces a novel statistical method of combining different enrichment tests into a single p-value, demonstrating both its value and shortcomings on simulated and real data. Our simulations show that GWASJET (the block jackknife joint GWAS enrichment test) has the highest statistical power of the investigated GWAS enrichment tests. This is not because it surpasses all other individual methods in power in all scenarios, but rather because in most scenarios it retains the power of the best-performing method, with few exceptions. However, it is computationally expensive, taking many hours of runtime in the single-core mode.

The primary limitation of the simulation study comes from the difficulty of investigating the full parameter space, given the high computational cost associated with the method. Many areas of the parameter space remain unexplored (low enrichment and very large sample size, low heritability but high enrichment, very high or very low heritability). We also assume 4,000 common variants are causal for the simulated phenotype in most studies, but in practice this number is unknown. It has been estimated that most traits have a target size (i.e. number of sites in the genome that would alter phenotype function, if a mutation occurred) in the millions to hundreds of millions [24], and there are 12,111 variants independently associated with height [25], so it is possible the number for many diseases may be substantially higher than 4000. Unfortunately, our simulation framework does not scale well beyond 5,000 causal variants, as the simGWAS method is limited in how many causal variants can lie within the same genomic region.

The application of this data to inflammatory bowel disease provided examples of the benefits of this approach, including an increase in the number of known important cell-types discovered, and a reduction in method-specific and seemingly spurious results (such as gametocytes). As expected, immune cell-types bear the overwhelming majority of IBD risk gene expression. This not only replicates prior long-standing laboratory and clinical studies in IBD [26], but also previous GWAS enrichment testing, which has shown clear signal within lymphoid (T-cells, B-cells) and myeloid (macrophages, dendritic cells and others) lineages [5]. While this does not exclude an important role for stromal and epithelial compartments in IBD aetiology, it certainly underscores the heavy role that immunity plays in this case.

We also replicate previous results showing that cell stimulation is frequently important for inducing IBD risk gene expression, particularly in myeloid cells [8]. In our analyses, the strongest myeloid enrichments were seen in macrophages stimulated with M-CSF, interferon gamma and Pam3Cys (a synthetic bacterial lipoprotein), monocytes stimulated using a blocking antibody to FcgammaRIIB, and dendritic cells stimulated with an agonist antibody to DC-SIGN. These molecules are known from previous literature to be important in regulation of immune processes [27–29]. However, it is important to note that the HPCA dataset these findings are based on is a collation of hundreds of studies from dozens of labs, and further validation is required to determine whether enrichment is truly down to the stimulation condition, rather than to confounders such as lab-specific differences in cell culture approach.

While eQTL studies of activated myeloid cells have been carried out before [30,31], they tend to use just one or two stimulatory molecules, often obvious candidates such as interferon gamma. Our results demonstrate that there are many different stimulations that may be important for cells to manifest IBD risk pathways, including many that are not top of researchers’ lists when designing experiments, but which may provide value as alternative stimulations in future eQTL studies. Having a focused but more comprehensive activation panel, possibly one that has been screened first using GWAS enrichment testing, could help us discover more context-dependent ways in which IBD risk loci act.

One result of this study, which we believe has not been reported before, is that in addition to being enriched for CD and UC risk genes, a wide range of immune cells are also enriched for variants that predict having CD compared to UC. In all cases these cell-types showed stronger enrichment of CD risk than UC risk, with the largest differences being seen in monocytes and gamma-delta T cells. This may represent CD having a higher heritability, and thus having a larger number of risk variants with larger effect sizes in the same shared IBD risk pathways. Finding ways of developing these results to control for global patterns of difference in effect size, and to identify UC-specific genetic risk pathways, would be of significant value to the field.

Further improvements could be made to the method, with the most important being to improve the computational speed. Running the test at scale requires a high-performance computing cluster, which narrows the appeal of the method to some researchers and imposes a carbon cost on research [32]. The major computational costs of the methods are in calculating LD scores and running SNPsea. Improvements to LD scoring are difficult, but as the LD scores for a given gene expression dataset are fixed, by generating and sharing pre-computed LD scores for the same gene expression dataset across different users the cost can be shared and reduced. Improvements to SNPsea, particularly to allow more pre-computation and to reduce the numbers of permutations required – for example, by using tail approximations [33] – would also reduce the computational cost substantially. Other improvements to the model, for example to introduce conditional testing to identify if enrichment of different cell-types are driven by the same genes, or modifying the method to analyze gene region data such as ATAC-Seq peaks, would allow the benefits of joint testing to be extended to other use cases.

## Materials and Methods

### Block jackknife GWAS joint enrichment test

#### Overview of the method

The aim of our method is to combine the evidence for GWAS enrichment, measured by their p-values, into a single p-value that is uniform under the null distribution. To do this, we take the simple approach of transforming each method’s one-tailed p-values into Z scores using the standard normal inverse cumulative distribution function, and then to sum the Z-scores together, and then transform this summed p-value into a final combined p-value.

Calculating this combined p-value requires us to know the distribution of this summed Z score under the null. A sum of standard normal Z-scores is distributed with an expectation of zero and an unknown variance which can be estimated from the correlation structure of the Z-scores. As the different enrichment tests are derived from the same summary statistics, we expect their Z-scores to be correlated, and estimate this correlation structure with a block jackknife approach, which is described in the following sections. We then use this to estimate the variance in the sum of Z-scores and then create a new combined Z-score that is standard normal under the null, which can be used to generate well calibrated P-values.

#### The distribution of the test statistic

Our test statistic is the sum of several correlated Z scores. We need to derive how this test statistic (from here onwards, named *S_bjk_*) is distributed. We will write the vector of test statistics for each of the *n* enrichment methods as the *Z* = *Z*_1_, *Z*_2_,…,*Z_n_* and assume that, under the null, this vector is drawn from a multivariate normal, *Z*∼*N*(0,*J_corr_*) where *J_corr_* is the correlation matrix of the Z-scores.

We can write the sum of the Z scores, 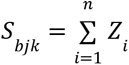, as vector product *S_bjk_* = (1, 1,…,1)·*Z^T^*

We use the affine transformation of a multivariate normal equation [34] and obtain

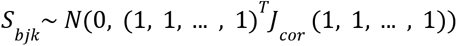

which, by abuse of notation, we will write as *S_bjk_* ∼N(0,∑*J_corr_*), where ∑*J_corr_* is the grand sum of the correlation matrix (the sum of all its elements).

Therefore, given a sum of dependent Z-scores, we can derive a joint Z-score by dividing the test statistic by the square root of the grand sum of the correlation matrix (i.e. by the standard deviation of *S_bjk_*), and use this joint Z-score to calculate a combined p-value of enrichment.

In practice, *J_corr_* is unknown and likely hard to derive analytically from the complex enrichment testing models, so we estimate it with a subsampling approach (the block jackknife).

#### Joint test algorithm

This section explains how the correlation between the Z scores is estimated first, and how the joint enrichment P-value is derived last.

The block jackknife is a data resampling method used to estimate the error variance and covariance of parameters estimated from data. It involves repeatedly removing parts of the data (“blocks”) and refitting parameters on the remaining data, and using the variance and covariance in these estimates to determine the error parameters. It first started to be widely used in genetics by the LDSC algorithm, which split the genome into many non-overlapping contiguous blocks of SNPs in order to calculate the variance on the heritability parameter estimated from summary statistics. We use the same approach here to calculate the covariance in the Z-scores across multiple methods.

We compute the whole-data estimates with all the *n* enrichment testing methods, and then recompute the estimates M times, while removing one block of SNPs from the analysis at a time. We will call an enrichment Z-score obtained after one block of SNPs is removed a “delete-value”, and the region of the genome that contains this block of SNPs the “delete-block”. We can then use the same equations as those described by the authors of LDSC [5,13,14] to compute our jackknife-derived covariance estimate:

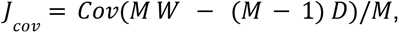

where M is the number of blocks, W is a row-matrix of whole data estimates (one column for each method, and each row filled with the same Z-scores computed on the whole data) and D is a matrix of of delete-values (*one* columns for each enrichment methods, and M rows containing the delete-value Z-score estimates for that block). We then obtain the correlations from the covariance structure

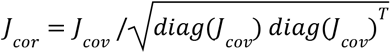

where diag(X) is the vector of the elements of the diagonal of matrix X.

Once we have this correlation structure, we can compute the result as a Z-score for each cell-type *c* as

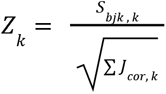

where *k* indexes cell-types, *S_bjk,k_* is the Z-score sum for the *k*-th cell-type, and ∑*J_corr,k_* is the sum of the elements of the correlation matrix *J_corr_* estimated for the *k*-th cell-type. These combined Z scores are transformed into enrichment p-values using the standard normal cumulative density function and adjusted for multiple testing to maintain a set false discovery rate using Benjamini-Hochberg.

#### Joint test implementation details

For our implementation of this joint test, we chose to use MAGMA (v.1.10), SNPsea (v.1.0.3) and LDSC (v.1.0.1) – all in binary setting – using default parameters for all. We used 200 equally sized blocks of SNPs for the block jackknife (the same number as used by LDSC).

For MAGMA, we first pre-compute all gene Z scores and calculate the whole data enrichment p-values, and then estimate MAGMA delete values for enrichment Z scores, we use the MAGMA parameter *--settings gene-exclude* to eliminate genes contained within the delete-block.

For LDSC and SNPsea, we calculate whole data Z scores using default settings, and then calculate delete values for each iteration of the block jackknife by analysing summary statistics where all SNPs contained within the deleted block have been removed. For SNPsea, genes from delete-blocks are also removed from the gene-set before testing.

We exclude methods from the combined p-value if the block jackknife estimate of the Z score variance for its 200-blocked estimates is zero, which is mostly an issue when SNPsea estimates are extremely significant (and thus always have the same, maximum, Z score).

### Simulation study of enrichment testing techniques

#### Overview of the simulation

We carry out simulation studies to assess the power of GWASJET compared to the three component tests (LDSC, MAGMA and SNPsea). We simulate a gene-set, then simulate GWAS summary statistics based on an underlying model with fixed parameters (sample size, heritability, enrichment of heritability within the gene-set, polygenicity, etc), and then apply our different enrichment tests to these simulated datasets to test the calibration (proportion of simulated null gene-sets that are found to be significant at p < 0.05) and power (number of truly enriched gene-sets that are found to be significant at p < 0.05).

#### Heritability modeling assumptions

In our simulation framework, we first select sets of causal variants and effect sizes under a simple polygenic heritability model, and then simulate GWAS summary statistics under a binary model using the R package simGWAS (v.0.2.0.1) [35].

In our simple model there are two categories of SNPs: ones inside (or close to, within a window of) a specific gene-set (referred to as gene-set SNPs), and ones that remain after excluding these (outside SNPs). These two categories of SNPs can have different densities of causal variants and different distributions of effect sizes. We write the density of causal variants (i.e. the probability that a randomly selected variant will be causal) as *p_c_*(where *c* is the index of the category, 1 for gene-set SNPs and 0 for outside SNPs). For a given causal variant indexed *i*, we assume that the effect size (log odds ratio) is drawn from a normal distribution *β*∼*N*(0,*σ_c_*^2^), where the variance parameter *σ_c_*^2^ is different for gene-set and outside SNPs.

To simulate a single GWAS, we simulate a gene-set by choosing 2000 genes uniformly at random, assigning each variant a value *c* based on its proximity to the nearest gene, sampling a set number of causal variants from each of the two SNP sets to give a causal density within each category of *p_c_*, sample a log odds ratio for each causal variant from *N*(0,*σ_c_*^2^), and then simulate genome-wide summary statistics for a given sample size and matrix of linkage disequilibrium between variants using simGWAS.

#### Converting from genetic architecture parameters to simulation model parameters

In our study, we simulated GWAS with different genetic architectures, defined based on the SNP heritability (i.e. trait variation explained by the SNPs, *h*^2^), the ratio of average heritability explained per SNPs inside and outside the gene-set, and the total number of causal variants in the genome. To do this, we need to translate from these genome-wide parameters to the parameters inputted into the simulation approach describe above.

We do this by noting that, under a linear model, the variance explained by the set of all SNPs in a given gene-set *c* is given by

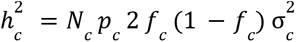

where *N_c_* is the total number of variants in the gene-set (and thus the number of causal variants in the gene-set is *N_c_p_c_*) and *f_c_* is the average allele frequency of the variants in the gene-set (which we assume to be equal to the average allele frequency of causal variants in the gene-set, as variants are selected independently of allele frequency in our simulations).

The three genetic architecture parameters are thus total heritability, given by 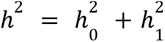, the number of causal variants, given by *N*_0_*p*_0_ + *N*_1_*p*_1_, and the ratio of heritability explained per SNP within and outside the gene-set, given by 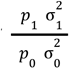. Once these parameters have been specified, and *f*_0_, *f*_1_, *N*_0_ and *N*_1_ have been estimated from the data, only one further free parameter needs to be set (either the ratio of causal variant densities, 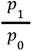, or the ratio of effect size variances, 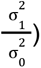. In this study, we make the additional assumption that 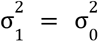 i.e. that effect sizes are the same in the two SNP sets and the two gene-sets only differ in their density of causal variants and the ratio of per-SNP heritability is the same as the ratio of causal SNP densities 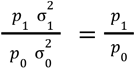. When this assumption is made, all simulation parameters can be uniquely determined by solving the above set of simultaneous equations given above.

Note that the above equations were derived under a linear model, but simulations are carried out under a binary logistic model, and thus the resulting GWAS will not have the desired heritability on the observed binary scale. We therefore run simulations at several simulated linear-scale heritabilities (with a large sample size of 1 million cases and controls to obtain narrow confidence intervals) and determine the observed heritability with LDSC, computing a calibration curve that can be used to select a linear-scale heritability parameter that gives the desired heritability on the observed binary scale.

#### Data and implementation details for simulations

The 2,000 genes per causal set are sampled from 22,350 NCBI RefSeq protein coding canonical transcripts on reference hg19 (build 37), downloaded from UCSC.

For most analyses we simulate 4,000 causal variants genome-wide, which are sampled from the HapMap3 [36] European SNP list, a set of well-imputed and well-behaving SNPs originally generated by the LDSC authors [13]. We chose to simulate 4,000 causal variants because GWAS of complex traits near saturation can achieve thousands (if not more) causal variants [25]. We tested causal SNP densities 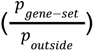 between 1 and 2, sample sizes from small GWAS (one thousand) to large biobank scale (one million individuals) with equal cases and controls, and three heritabilities (10, 20 and 30%, representing a low-heritability), proximity window choice (LDSC model uses 100 kilobases upstream and downstream, whereas MAGMA uses 10 and 1.5 respectively) and polygenicity level (4,000 total causal variants, for a highly polygenic trait, or 200, for a less polygenic trait). For all of these, the trait h^2^ is set at 20% (except for the heritability comparison simulations), the sample size of the GWAS is set to 65 thousand cases and controls (except for the sample size comparison simulations), and the ratio of causal SNP density is set to 1.75 (except for the simulation that compares ratios of causal SNP densities and the SNP-to-gene window model simulations). The SNP-to-gene window model simulations were altered to have a lower causal density ratio (1.25 instead of 1.75) in order to match the number of “outside gene-set” SNPs between the LDSC and MAGMA window simulations.

For each simulation, simGWAS was run separately within 1703 approximately-independent LD blocks in the human genome obtained from the literature [37] and these blocks were then combined to produce genome-wide GWAS summary statistics as suggested by the simGWAS documentation. Allele frequencies and linkage disequilibrium, required to run simGWAS, were calculated using European samples of the 1000 Genomes data [38].

We estimate the type 1 error rate using the null model of a causal SNP density of 1, and the power across all other parameter values. We carry out 300 replications for each simulation, giving reasonable confidence-intervals on the estimates without requiring excessive runtimes. The p-value threshold (ɑ) used to assess power and type 1 error in all simulations is 5%. Clopper-Pearson 95% exact confidence intervals for the statistical power were computed with the PropCIs (v.0.3) R package.

### Application of the method to inflammatory bowel disease summary statistics

To test GWASJET with real datasets, we focused on inflammatory bowel disease risk genes in disease-relevant cell-types. This involved collating a large set of publicly-available gene expression datasets (Table 2), including general maps of human cells (Human Primary Cell Atlas, single-cell blood cell data from 10X), immune-focused datasets (Human Cell Atlas immune census) and single-cell data from intestinal biopsies (from IBD patients and healthy controls). We tested for enrichment of risk genes in these cell-types using summary statistics for both IBD subtypes, CD and UC, as well as a novel analysis of the genetic differences between CD and UC patients using synthetic genome-wide CD vs UC GWAS summary statistics.

**Table 2:**
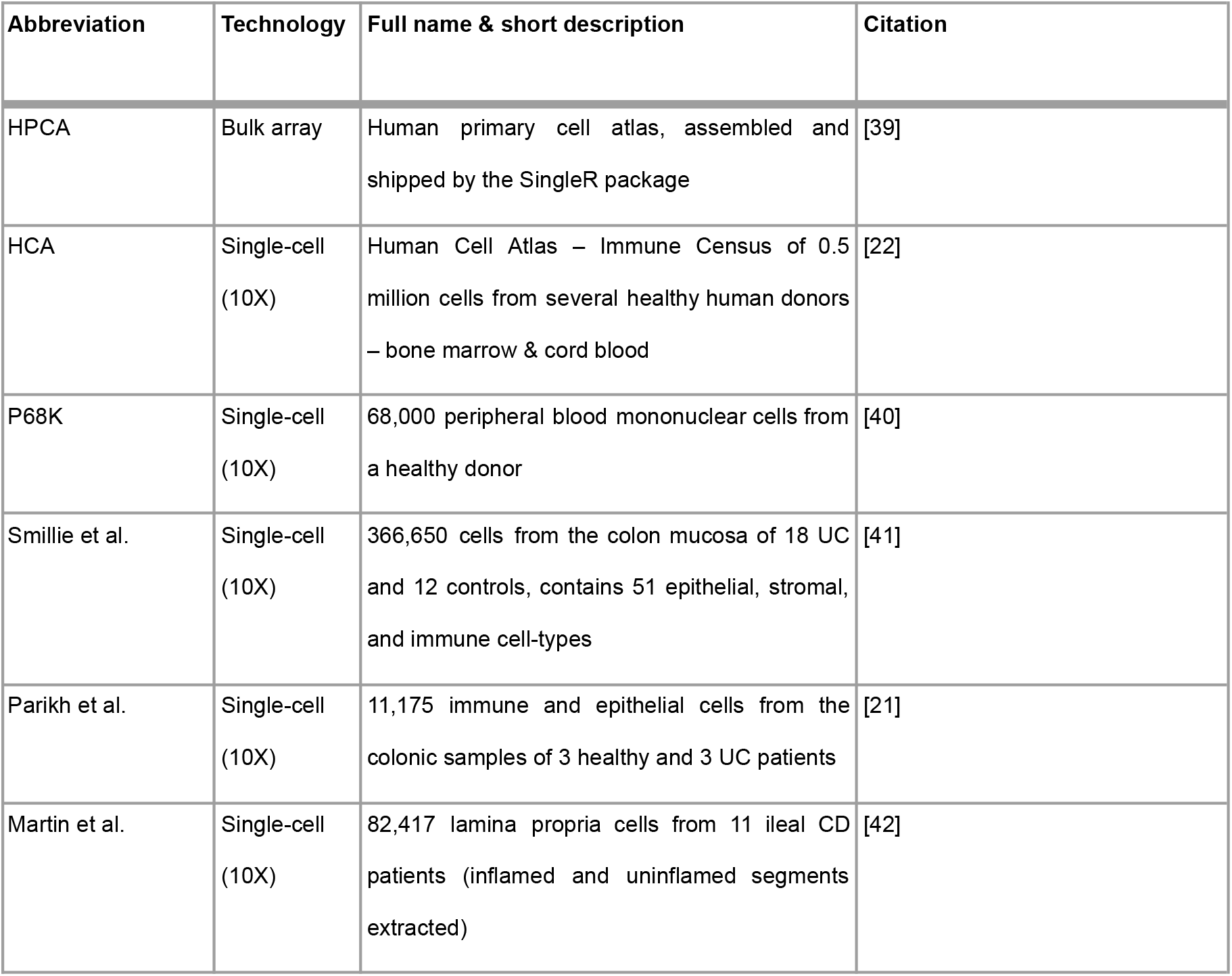
Expression datasets used in this study.

| Abbreviation | Technology | Full name & short description | Citation |
| --- | --- | --- | --- |
| HPCA | Bulk array | Human primary cell atlas, assembled and shipped by the SingleR package | [39] |
| HCA | Single-cell<br>(10X) | Human Cell Atlas – Immune Census of 0.5 million cells from several healthy human donors – bone marrow & cord blood | [22] |
| P68K | Single-cell<br>(10X) | 68,000 peripheral blood mononuclear cells from a healthy donor | [40] |
| Smillie et al. | Single-cell<br>(10X) | 366,650 cells from the colon mucosa of 18 UC and 12 controls, contains 51 epithelial, stromal, and immune cell-types | [41] |
| Parikh et al. | Single-cell<br>(10X) | 11,175 immune and epithelial cells from the colonic samples of 3 healthy and 3 UC patients | [21] |
| Martin et al. | Single-cell<br>(10X) | 82,417 lamina propria cells from 11 ileal CD patients (inflamed and uninflamed segments extracted) | [42] |

#### Pre-processing and clustering of expression data

Single-cell expression datasets were library-normalised (counts per 10,000) for each cell and log(1+X) transformed. When filtered datasets were provided by the original studies, we used those, otherwise for unfiltered datasets (such as the Human Cell Atlas immune census) we filtered out cells that had fewer than 500 unique molecular identifiers (UMIs), number of genes detected per cell > 5000 (to remove doublets), percentage of gene expression of mitochondrial origin (perc.mt.) > 10%, and we regressed out the mitochondrial percentage (perc.mt). All these analyses were carried out in Seurat (version 4.0.3) [43]. To provide comparable cell-type labels for cells across single-cell datasets, we used SingleR (version 1.6.1) [44] with the Human Primary Cell Atlas (HPCA) panel of markers supplied by the package to assign each cell to a cell-type at both L1 (broad cell-type) and L2 (fine cell-type) level.

Once pre-processed and clustered, gene specificity scores for each cell-type were derived using two methods previously described in the literature for this purpose: the Expression Weighted Celtype Expression score (EWCE, v.1.0) [19] and the equal-variance t statistic (tstat) [5]. The first is based only on mean expression, whereas the second uses both mean and variance to score genes. These scores were used both as continuous predictors, and were also thresholded to make cell-type specific gene-sets by selecting genes in the top 10% of each cell specificity score.

To aid visualization, we also computed a hierarchical clustering tree of gene expression for each dataset based on the Euclidean distances between the cell-types’ gene expressions, and plotted next to the ordered cell-types with the ggdendro 0.1.22 [45].

#### IBD GWAS summary statistics

Summary statistics (in hg19 coordinates) for IBD, CD and UC, were obtained from the latest published international GWAS meta-analysis study [46]. Part of the munging into the right summary statistics file formats (which differ between MAGMA, LDSC and SNPsea) was done using MungeSumstats Bioconductor package [47], and also with functions inspired from its codebase.

To generate post-hoc genome-wide log odds ratio estimates, standard errors and p-values for a CD vs UC GWAS (i.e a case-control study where CD patients are cases and UC patients are controls), we use a technique for carrying out meta-analysis with overlapping samples [48]. In our case, this gives the following formulas that we applies to each individual SNP:

β*_CD versus UC_* = β*_CD_* – β*_UC_*, where β is the log odds ratio of a SNP in the summary statistics.

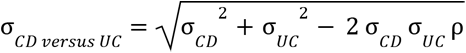,where σ is the standard error of the SNP and 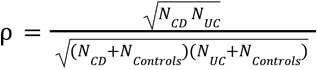 is the correlation in errors for case-control GWAS with overlapping controls, from equation 7 in [48]. In this case, ρ=0.249.

We then calculate a Z score and p-value:

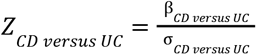

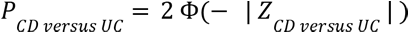, where Φ is the standard normal cumulative distribution function.

#### Enrichment testing

We applied GWASJET to each of the datasets above. We used 10 component tests in the joint test, using combinations of three different tools (LDSC, SNPsea, MAGMA), the two different ways of scoring cell-type gene specify discussed above (EWCE and tstat) and two ways of using these specificity scores (discrete top 10% gene-sets, “top10%”, or as linear continuous scores, “linear”). This sums to 10 rather than 12, as only MAGMA and SNPsea have currently supported modes that can analyse linear predictors. We combined these 10 tests into a single combined score using the block jackknife joint GWAS enrichment test in the same way as described for the simulation results, and p-values were corrected for multiple testing using Benjamini-Hochberg to maintain a false discovery rate of 5%.

#### Gene ontology enrichment testing

In order to test whether cell-type-specific genes that are also IBD risk genes, in particular, are overrepresented in specific pathways we perform the following analysis: We use MAGMA to obtain gene-to-trait association P-values, producing 1191 Benjamini Hochberg-significant genes for IBD. Separately, for each cell-type, the top 2,000 genes by expression specificity (t-statistic) were selected as markers. Then, the risk genes that were in the top 2,000 markers for that cell-type were tested for enrichment in Gene Ontology (GO) terms, when compared to a “universe” parameter (otherwise called the background) consisting of all the 2,000 markers. This was done using the enrichGO function of the clusterProfiler v.4.0.0 R package and its builtin gene ontology lists, using a Pval cutoff of 0.01 and a Qval cutoff of 0.05 and looking at all the 3 ontologies (BP, CC and MF).

## Code and data availability

An R package, entitled GWASCellTyper, with functions to apply the GWASJET method, as well as to functions to help running MAGMA, LDSC and SNPsea and process their input and output data is available at this link: https://github.com/alexandruioanvoda/gwascelltyper

All data used in this study is publicly available and cited throughout the methods section.

## Acknowledgements

The authors would like to acknowledge a number of scientists, including Prof. Alison Simmons, Dr. Agne Antanaviciute, Dr. Stephen Sansom, Dr. Kevin Rue-Albrecht, Dr. Chris Eijsbouts, Dr. Christiaan de Leeuw and Dr. Nathan Skene for valuable feedback and insightful discussions. Analyses were carried out using the Oxford Biomedical Research Computing (BMRC) facility, a joint development between the Wellcome Centre for Human Genetics and the Big Data Institute supported by Health Data Research UK and the NIHR Oxford Biomedical Research Centre. The views expressed are those of the author(s) and not necessarily those of the NHS, the NIHR or the Department of Health. This research was funded in whole, or in part, by the Wellcome Trust (208750/Z/17/Z). For the purpose of Open Access, the author has applied a CC BY public copyright licence to any Author Accepted Manuscript version arising from this submission.

---

**Supplementary figure 1:**
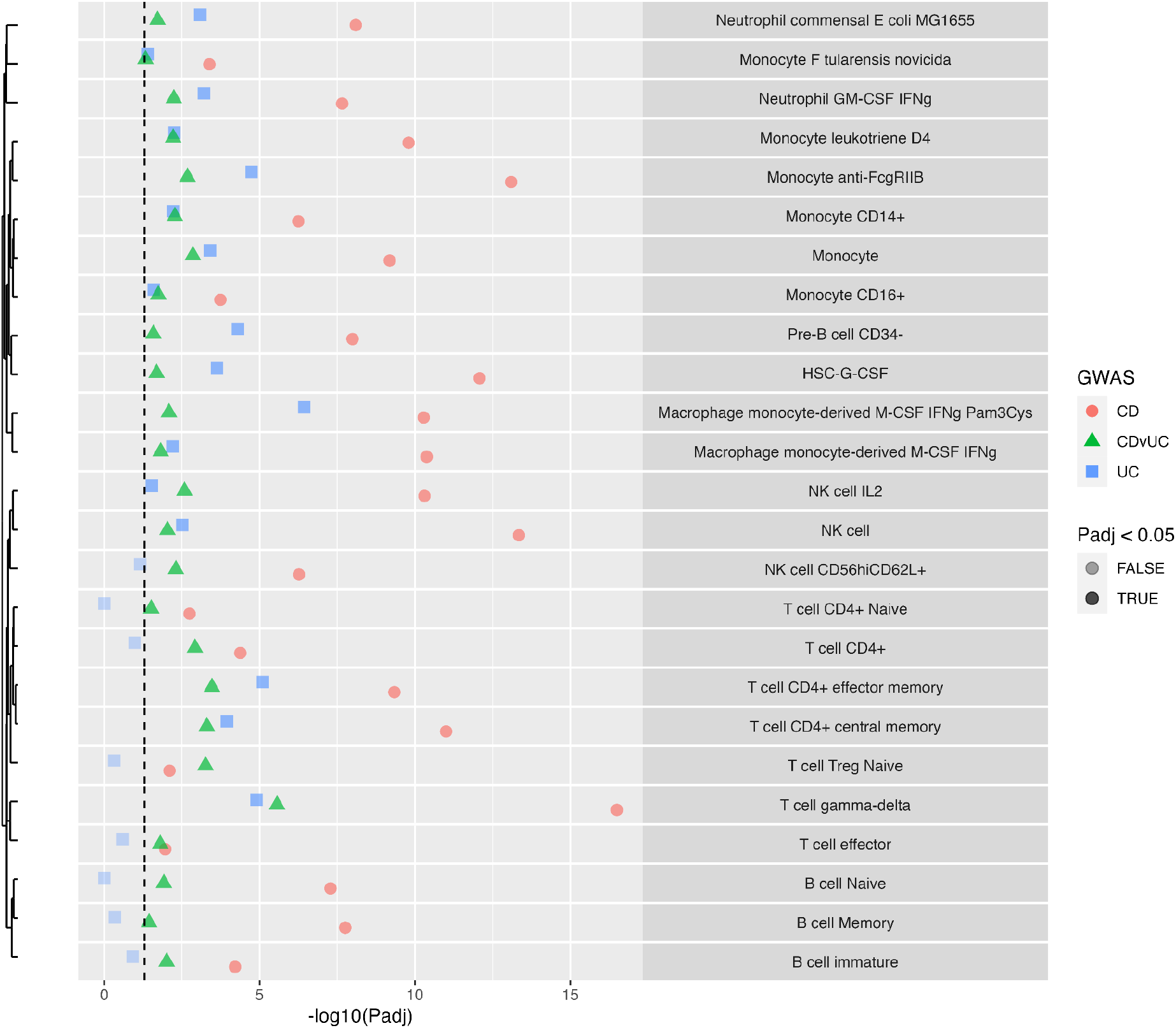
Differences in enrichment for disease subtype loci within finer-grained cell-types. Enrichment of risk loci for cell-type specific gene expression for CD and UC, along with enrichment for loci that differ between CD and UC (CDvUC), showing only the significant fine-grained cell-types from HPCA. The dashed line shows adjusted p < 0.05.

